# *Diutina (Candida) rugosa* complex: The biofilm ultrastructure, extracellular matrix, cell wall component and antifungal susceptibility to amphotericin B, caspofungin, fluconazole and voriconazole

**DOI:** 10.1101/2020.03.30.016246

**Authors:** Sri Raja Rajeswari Mahalingam, Priya Madhavan, Chong Pei Pei

## Abstract

The genus *Candida* is the most common etiological factor of opportunistic fungal infections in humans. The virulence of *Candida* species is due to a wide repertoire of factors, specifically, the ability to form biofilms. Medical devices such as intravenous catheters, prosthetic heart valves and surgical interventions provide pathogenic microorganisms with a surface to adhere to form biofilm. The objectives of this study were to investigate the biofilm ultrastructure of *Diutina* (*Candida) rugosa* (*D. rugosa*) at different developmental phases using Confocal scanning laser microscopy (CSLM) and scanning electron microscopy (SEM), quantify β-glucan, total carbohydrate and total protein in the extracellular matrix (ECM) using enzymatic β-glucan kit, phenol-sulfuric acid method and Bradford’s method, respectively, and to identify Sessile Minimum Inhibition Concentrations (SMICs) of amphotericin B, caspofungin, fluconazole, and voriconazole using serial doubling dilution. From the SEM micrographs, *D. rugosa* biofilms were composed of adherent yeast cells and blastospores with hyphal elements. The ultrastructure of the yeast cells was collapsed and disfigured upon exposure to amphotericin B, fluconazole and voriconazole and the biofilms presented with punctured yeast morphology upon exposure to caspofungin at their respective SMICs. The matrix thickness of embedded yeast cells from CLSM micrographs was 3.9µm at 48h. However, there was reduction in the thickness of the biofilms upon antifungal exposure. The antifungal exposed biofilms exhibit bright, diffuse, green-yellow fluorescence that were not seen in the control. *D. rugosa* biofilm matrices revealed 172.57µg/mL of carbohydrate, and 27.11µg/mL of protein content. The β-glucan yield in *D. rugosa* complex planktonic cells were in the range of 2.5 to 4.38%, on the contrary, β-glucan was not detected in the ECM. The SMICs of *Diutina* biofilm for amphotericin B is 1024μg/mL, caspofungin is 512 μg/mL, whereas fluconazole and voriconazole is 2048 μg/mL, respectively.

## Introduction

The yeast *Candida* is ubiquitous in nature. It is a commensal microorganism that asymptomatically colonizes the mucocutaneous system of the human body including skin and its appendages, respiratory, gastrointestinal and genitourinary tracts [1, 2]. Many species of *Candida* are opportunistic fungal pathogens. Candidiasis is the most frequent in immunocompromised hosts due to malignancy, those under cytotoxic immunosuppressants and patients with HIV infection [3]. However, it is also notable among critically ill, prolonged duration in ICU stay and non-immunocompromised patients [1].

*Candida rugosa (C. rugosa)* is a rare cause of candidemia but have been described as an emerging fungal pathogen [4-6]. Although rare, it is noteworthy to recognize due to its resistance to antifungal agents particularly to amphotericin B and azole drugs [4, 5, 7]. Phylogenetically, *C. rugosa* is considered as a species complex that comprises four taxa., i.e *C. rugosa, C. pseudorugosa, C. neorugosa and C. mesorugosa* [8]. In recent past, phylogeny of *C. rugosa* complex was reassessed and this complex was relocated into a new genus *Diutina* [9, 10]. Kunnamwong and co-workers have identified the presence of a clade that is not associated with any known teleomorphic genus based on phylogenetic analysis of small and large rRNA gene subunits of three strains isolated from rice leaf tissue. Along with these strains, seven other *Candida* species have been affiliated with this clade including *C. rugosa, C. pseudorugosa, C. neorugosa and C. mesorugosa*. These strains were later relocated into this clade under a newly proposed genus *Diutina* [10]. *Diutina rugosa* (*D. rugosa*) was referred as *C. rugosa* in earlier studies and was classified under genus *Candida*, the literature on the latter will be used to describe the pathogenicity of *D. rugosa. C. rugosa* was ranked the ninth most common of 16 *Candida* species isolated from clinical specimens in the ARTEMIS DISK antifungal susceptibility program over the period 1997-2005. Over that period, the antifungal susceptibility test showed a decrease in the susceptibility towards antifungal agents [11]. *C. rugosa* is highly prevalent in fungemia with invasive procedures. The common clinical features of *C. rugosa* infections are intravenous catheter-associated candidemia and frequent colonizer in burned patients [1, 4, 5].

Pathogenicity of *Candida* species are attributed with many virulence factors – dimorphism, biofilm formation and secretion of hydrolytic enzymes. The ability of *Candida* species to form biofilm is the major factor of therapeutic failure due to their intrinsic tolerance to antifungal agents [12]. Biofilm provides a survival advantage to *Candida* species by providing protection from host and hostile environments [12]. Candidemia associated to biofilm forming isolates have higher mortality rates than non-biofilm forming isolates [13]. The high prevalence of *D. rugosa* infections in intravenous catheter could be associated to biofilm formation. *Candida* species such as *C. albicans, C. parapsilosis, C. tropicalis, C. glabrata, C. dubliniensis*, and *C. krusei* are all capable of forming biofilm [2, 14]. However, the biofilm structures vary among the *Candida* species. It has been noted that the extracellular matrix consists of contrasting amounts of carbohydrates and proteins for different *Candida* species [2].

Notably, *Candida* species biofilm is up to 1000-fold more resistant to antifungal agents than their planktonic counterparts [15, 16]. Studies has shown that *C. rugosa* is notable for decreased susceptibility to azoles and amphotericin [7, 8, 17]. The biofilm of six *C. rugosa* isolates from blood, urine, vaginal swap and bronchoalveolar has increased resistance to amphotericin B (MIC ≥ 8µg/mL) and fluconazole (MIC ≥ 128 µg/mL) as compared to planktonic cells [7]. In Malaysia, a recent study has reported that the MICs *C. rugosa* complex planktonic cells isolated from blood ranged from 1.8 to 2 μg/mL, 2 to 5 μg/mL, 7 to 48 μg/mL and 0.125 to 0.5 μg/mL for amphotericin B, caspofungin, fluconazole and voriconazole, respectively [18]. Interestingly, in an outbreak of candidemia caused by *C. rugosa* in Brazil between 1995 to 1996, all the *C. rugosa* isolates were susceptible to fluconazole (MIC range 0.125 to 0.5 µg/mL) and amphotericin B (MIC range 0.5 to 1 µg/mL) [19]. Similarly, in Kolkata, India, between 2011 to 2015, two out of 70 samples isolated blood culture were positive for *C. rugosa* and both the isolates were susceptible to amphotericin B, fluconazole, voriconazole but resistant towards itraconazole [20]. Indeed, there are many controversies about the antifungal susceptibility patterns of *D. rugosa* complex. The diversity of antifungal MIC’s within the strains of *D. rugosa* complex may be associated to geographic variations, site of infections and inter- and intraspecies variations [8].

In Malaysia, *D. rugosa* is considered a rare but an emerging fungal pathogen. In a prospective laboratory-based surveillance conducted at University of Malaya Medical Centre, Malaysia between year 2000 and 2013, *C. rugosa* was ranked sixth with 78 out of 34 392 total (0.23%) isolates from blood, oral cavity and urine samples [21]. In a retrospective study conducted at Hospital University Science Malaysia, between year 2010 and 2014, two out of 134 (1.5%) isolates from blood samples were identified as *C. rugosa* [22]. Similarly, in a study conducted at another teaching hospital in Kuala Lumpur, Malaysia between year January and July 2009, one out of 82 (1.2%) isolates from blood fluids was positive for *C. rugosa* [23].

The paradigm shift from common to rare but emerging *Candida* species infections manifest a concern in the reduced susceptibility to antifungal agents and its clinical relevance. Therefore, the aim of this study was to investigate the ultrastructure, bio-chemical profile: β-glucan, carbohydrate and protein contents of *D. rugosa* planktonic cells and biofilm, in addition to its antifungal susceptibility towards amphotericin B, caspofungin, fluconazole and voriconazole.

## Materials and methods

### Collection of *D. rugosa* complex and culture conditions

A total of seven *D. rugosa* strains as pure cultures were obtained from two tertiary referral hospital laboratories in Malaysia. The strains were isolated between year 2007 and 2016 from blood samples and kept in pure cultures at −80°C in the respective hospital laboratories. In our laboratory, all the strains were thawed and sub-cultured on Sabouraud Dextrose Agar (SDA) (Oxoid, Hampshire, UK) to obtain pure cultures and stored in 15% (vol/vol) glycerol (Fisher Scientific UK Ltd., England) at −80°C until further used. One reference strain from the American type Culture Collection (ATCC) of *D. rugosa* ATCC 10571 was used as a positive control in this study.

### Identification of *D. rugosa* Complex

#### CHROMagar Candida

CHROMAgar™ Candida (CHROMagar, Paris, France) was prepared according to the manufacturer’s guidelines. The isolates were inoculated onto the CHROMAgar™ plates and incubated at 37°C for 48 h. The appearance of colonies, including color, size, and textures on CHROMAgar™ Candida were observed. Small colonies of brilliant blue with pale border on the CHROMAgar™ plate indicate the presence of *D. rugosa*. Biochemical tests and molecular identification are then carried out for further confirmation

#### Biochemical tests

Biochemical tests were performed using RapID Yeast Plus System (Remel, USA) following the manufacturer’s guidelines.

#### Molecular identification

The DNA of the isolates were extracted using GeneAll DNA extraction kit (GeneAll Biotechnology, Korea). The primers used were ITS1 and ITS4, which amplify the internal transcribed spacer regions (ITS) of the ribosomal DNA. The amplicons were sequenced in both directions using primer pair ITS1 (5’-TCCGTAGGTGAACCTGCGG-3’) and ITS4 (5’-TCCTCCGCTTATTGATATGC-3’). The sequences of each isolates were compared to the NCBI GenBank database using BLAST tool.

### Quantification of β-glucan

The procedures for β-glucan measurement in the cell wall planktonic cells and ECM of biofilm were performed using enzymatic yeast β-glucan kit purchased from Megazyme International (Ireland), following the manufacturer’s guidelines.

### Antifungal susceptibility testing of biofilm

The antifungal susceptibility test for sessile (biofilm) cells was performed using broth serial doubling dilution method according to previously described protocol [24]. The biofilm was tested against four different antifungal agents which were amphotericin B (Nacalai Tesque, Inc., Japan), caspofungin (Sigma Aldrich, USA), fluconazole (Liofilchem, Italy), and voriconazole (Liofilchem, Italy). All antifungal agents were prepared using dimethyl sulfoxide (DMSO) (R & M Chemicals) as solvent and RPMI 1640 (Sigma Aldrich, USA) as diluent. The biofilm was grown over 48 hours according to previously described protocol [25], using 96-well microtiter plates. It was washed thrice with PBS buffer before the addition of the antifungal agents and incubated at 37°C for further 48 h. Concentrations in the range of 2048-0.125 µg/mL were used for all the tested antifungal agents. After the incubation period, the plates were washed thrice with PBS and XTT reduction assay was performed as described in the preceding section. The tests were carried out on three different occasions in quadruplicate and the average was calculated.

### Antifungal susceptibility testing of planktonic cells

The antifungal susceptibility test to determine minimum inhibitory concentration (MIC) of planktonic cells was performed using broth microdilution method according to the Clinical & Laboratory Standards Institute, M27Ed4 [26]. The planktonic cells assays carried out on three different occasions in quadruplicate.

### Matrix composition of biofilm

For the analysis of matrix composition of biofilm, total carbohydrate and total protein were performed. The biofilm growth condition and extracellular matrix (ECM) extraction were adapted from previously described protocols [25]. Total carbohydrate was estimated according previously described protocols with a slight modification, using glucose as a standard [27]. Total protein was estimated according to previously described protocol using bovine serum albumin (BSA) [28]. *C. albicans* ATCC 90028 was used as a positive control for this study. The assay was carried out in triplicate on three different occasions. The ECM supernatant was stored at −20°C until used for FT-IR spectral measurement.

### Fourier Transform Infrared Spectroscopy (FT-IR)

The spectra were recorded with FTIR-Fourier Transform Infrared Spectroscopy (Spectrum 100 FT-IR, Perkin Elmer, UK) ranging between wave numbers 4000 and 400 cm^−1^. Spectra from glucose and BSA were used as reference to compare the spectra from samples.

### Specimen Preparation for Confocal Laser Scanning Microscopy (CLSM)

The sample preparation for CLSM imaging was performed according to previously described protocol with slight modifications [29]. The biofilm was grown using 2-well cell culture chamber slide (SPL Life Sciences Co., Ltd., Korea). For the control (untreated) samples, 1mL of the standardised cell suspension (1×10^6^ cells/mL) were pipetted into each chamber of the 2-well cell culture chamber slide and incubated at 37°C at 1.5, 6, 12, 24, 48 and 72 h, respectively. For the treated samples, 48 h biofilm were grown as described above using 2-well cell culture chamber slide and then treated with antifungal drugs for a further 48 h. The concentration of antifungal agents was selected based on the SMIC values from the antifungal susceptibility test for all four antifungal agents performed in the preceding section. Following the incubation period, the chambers were washed twice with 1 mL sterile saline before staining with Live/Dead Yeast Viability stain (Invitrogen, Thermo Scientific, USA). The Live/Dead Yeast viability stain was prepared according to the manufacturer’s guidelines. The images were captured on a CLSM (TCS SP5 II, Leica) using LAS software. Microscopy was performed in duplicate on two different occasions.

### Specimen Preparation for Scanning Electron Microscopy (SEM)

The sample preparation for SEM imaging was performed according to previously described protocol with slight modifications using 22mm sterile cell culture coverslips (ThermoFisher Scientific, USA) [30]. The coverslips were carefully placed on the bottom of a 6-well plate. Then, 3 mL of the standardised cell suspension (1×10^6^ cells/mL) were pipetted into each well of the 6-well plates and incubated at 37°C at 1.5, 6, 12, 24, 48 and 72 h, respectively. After the incubation period, the wells were washed twice with sterile saline. The samples were then fixed with 2.5% cacodylate buffered glutaraldehyde (0.2 M; pH 7.2) (Sigma-Aldrich, USA) for 5 h at 4°C. For the treated samples, 3 mL of the standardised cell suspension (1×10^6^ cells/mL) were pipetted into each well of the 6-well plates containing coverslip and incubated for 48 h at 37°C before treating with antifungal agents for a further 48 h. The antifungal agent concentration was selected based on the SMIC values from the antifungal susceptibility test for all four antifungal agents performed in the preceding section. The samples were then processed according to standard protocol for SEM and the topographic features of the biofilm were visualised with JEOL JSM-6400 Scanning Electron Microscope.

## Statistical analysis

Results were analysed using SPSS software (IBM SPSS Statistics for Windows, Version 23.0). Values were expressed as the means ± standard deviation. The OD values from individual *Candida* biofilm were compared by student’s *t* test. Analysis of variance (ANOVA) with an associated *post hoc* Tukey’s test was used for multiple-comparisons of individual *Candida* biofilm. All tests were performed with a confidence level of 95%.

## Results

### Identification of *D. rugosa* complex

Originally, the identification of the species was performed using CHROMagar™ Candida medium. All the isolates including seven *C. rugosa* and the reference strains, *D. rugosa* ATCC 10571 produce small colonies of brilliant blue with pale border, respectively. From the biochemical analysis using RapID Yeast Plus System, all the isolates were identified as *C. rugosa*. From the ITS sequence BLAST search using GenBank analysis, six isolates were identified as *D. mesorugosa* with 100% similarity (GenBank accession number HM641831). One isolate, was identified as *D. rugosa* with 100% similarity (GenBank accession number HM641832). Therefore, *D. rugosa* isolate was designated as Dr1 1, Dm2, Dm3, Dm4, Dm5 and Dm6 as *D. mesorugosa*.

### Analysis of β-glucan content

In this study, one *D. rugosa* and six *D. mesorugosa* clinical isolates, and three ATCC reference strains: *C. albicans* ATCC90028, *C. parapsilosis* ATCC22019 and *D. rugosa* ATCC 10571 were tested. Results in Fig 1 show that the β-glucan content in the cell wall of *Diutina* planktonic cells and three reference strains, *D. rugosa* (ATCC 10571), *C. albicans* (ATCC 90028), and *C. parapsilosis* (ATCC 22019). *D. mesorugosa* (Dm1) has the highest concentration of β-glucan with 4.38% and *C. albicans* (ATCC 90028) and *C. parapsilosis* (ATCC 22019) have the lowest concentration of β-glucan with 1.77%, respectively. The yield of the β-glucan in *D. rugosa* (Dr1) was 4.26%, while that in the reference strain *D. rugosa* (ATCC 10571) was only 1.77%. Results from the β-glucan content of *D. rugosa* and *D. mesorugosa* planktonic cells, showed statistically significant difference between the *Diutina* complex (t-test, p = 0.245).

**Fig 1.**
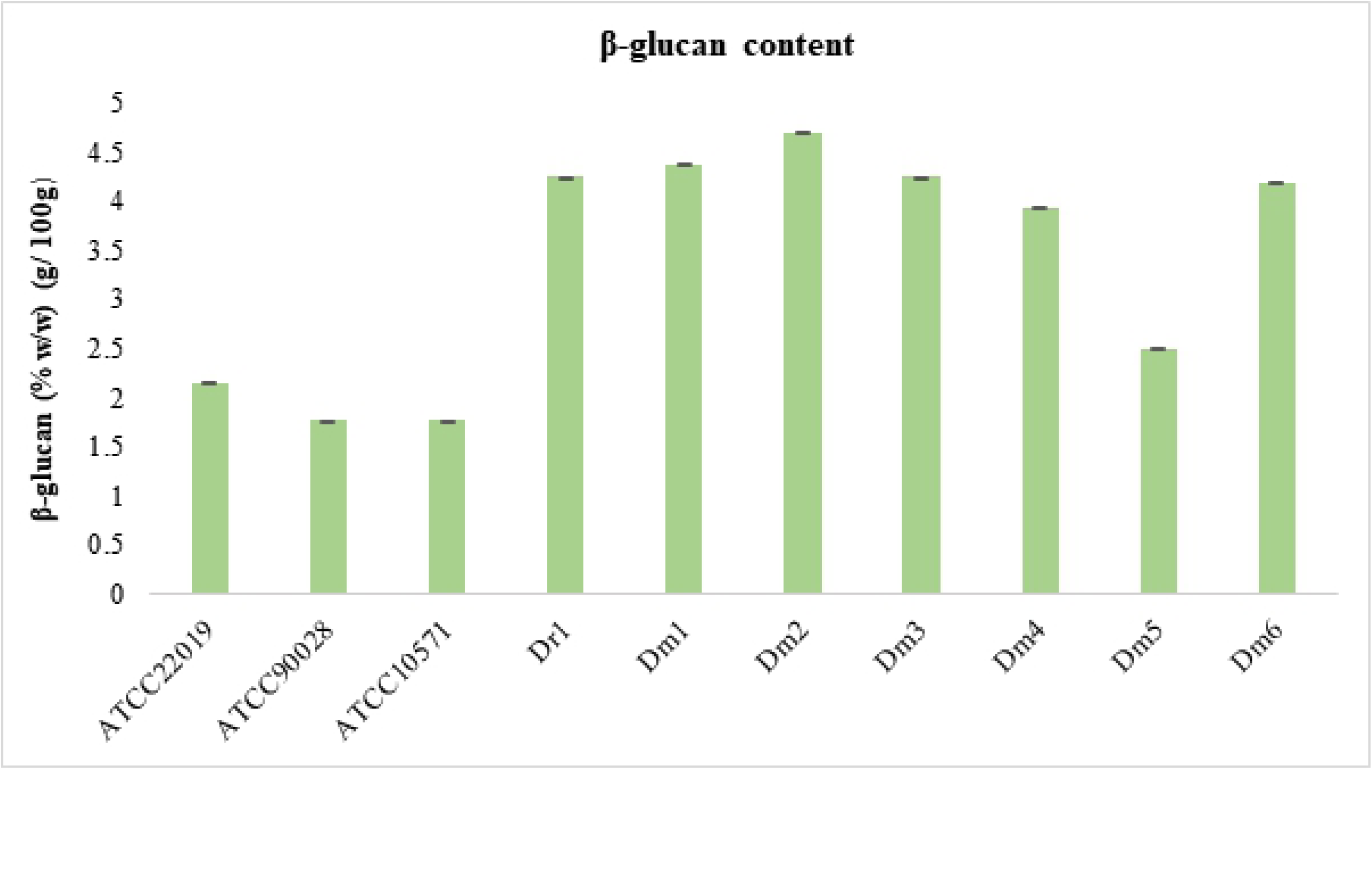
Analysis of β-glucan content in the cell wall of *D. rugosa* complex planktonic cells.

Dr1 *(D. rugosa)*, Dm1, Dm2, Dm3, Dm4, Dm5 and Dm6 (*D. mesorugosa), D. rugosa* ATCC 10571, *C. albicans* ATCC 90028, and *C. parapsilosis* ATCC22019. Error bars represent standard deviation (n=3).

### Analysis of Extracellular Matrix of biofilm

The extracellular matrix (ECM) were analysed for total carbohydrate and total protein by using colorimetric methods. In this study, one *D. rugosa* and six *D. mesorugosa* clinical isolates, and three ATCC reference strains, i.e. *C. albicans* ATCC90028, *C. parapsilosis* ATCC22019 and *D. rugosa* ATCC 10571 were tested. Results in Fig 2 show that the ECM components in all the biofilm are primarily composed of carbohydrate with a lower concentration of protein. Interestingly, *C. albicans* has the lowest total carbohydrate content as compared to all the *Diutina rugosa* biofilm. The total carbohydrate of *Diutina* complex ranged from 172.57 to 860.67 µg/mL of carbohydrate whereas total protein ranged from 0.307 to 0.414 µg/mL. Results from the total protein content of *D. rugosa* and *D. mesorugosa* biofilm, showed no statistically significant difference between the species of *Diutina* complex (t-test, p = 0.165). However, results from the total carbohydrate content of *D. rugosa* and *D. mesorugosa* biofilm, showed statistically significant difference between the *Diutina* complex (t-test, p = 0.04).

**Fig 2.**
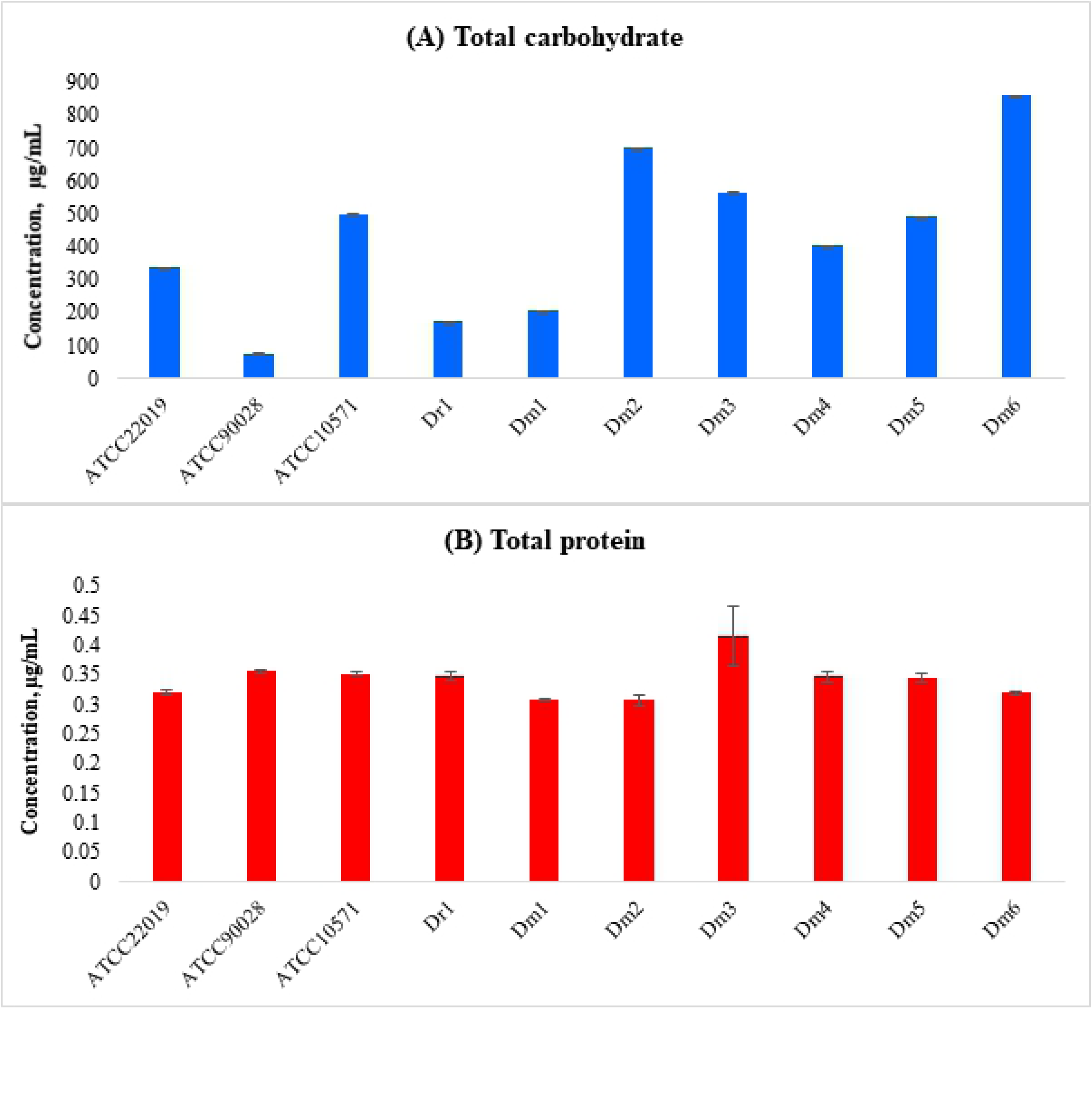
Analysis of ECM content of *D. rugosa* complex biofilm.

(A) total carbohydrate and (B) total protein from ECM components extracted from biofilm of Dr1 *(D. rugosa)*, Dm1, Dm2, Dm3, Dm4, Dm5 and Dm6 (*D. mesorugosa), D. rugosa* ATCC 10571, *C. albicans* ATCC 90028, and *C. parapsilosis* ATCC22019. Error bars represent standard deviation (*n=3*).

To obtain a better knowledge of the ECM extracted from *Diutina* biofilm, characterisation was performed using Fourier-transform infrared spectroscopy (FTIR) technique. Fig 3 shows the FTIR spectra of ECM produced by *Diutina* biofilm. From the FTIR spectral profile, absorption near 1641 cm^−1^ arises from the C=O stretching and absorption near 1551 cm^−1^ arises from N-H bending. Both the aforementioned absorption indicates the amide link in a protein molecule. The medium absorption at 3269 cm^−1^ indicates the O-H bond of glucose in the EPS components in *Diutina* biofilm.

**Fig 3:**
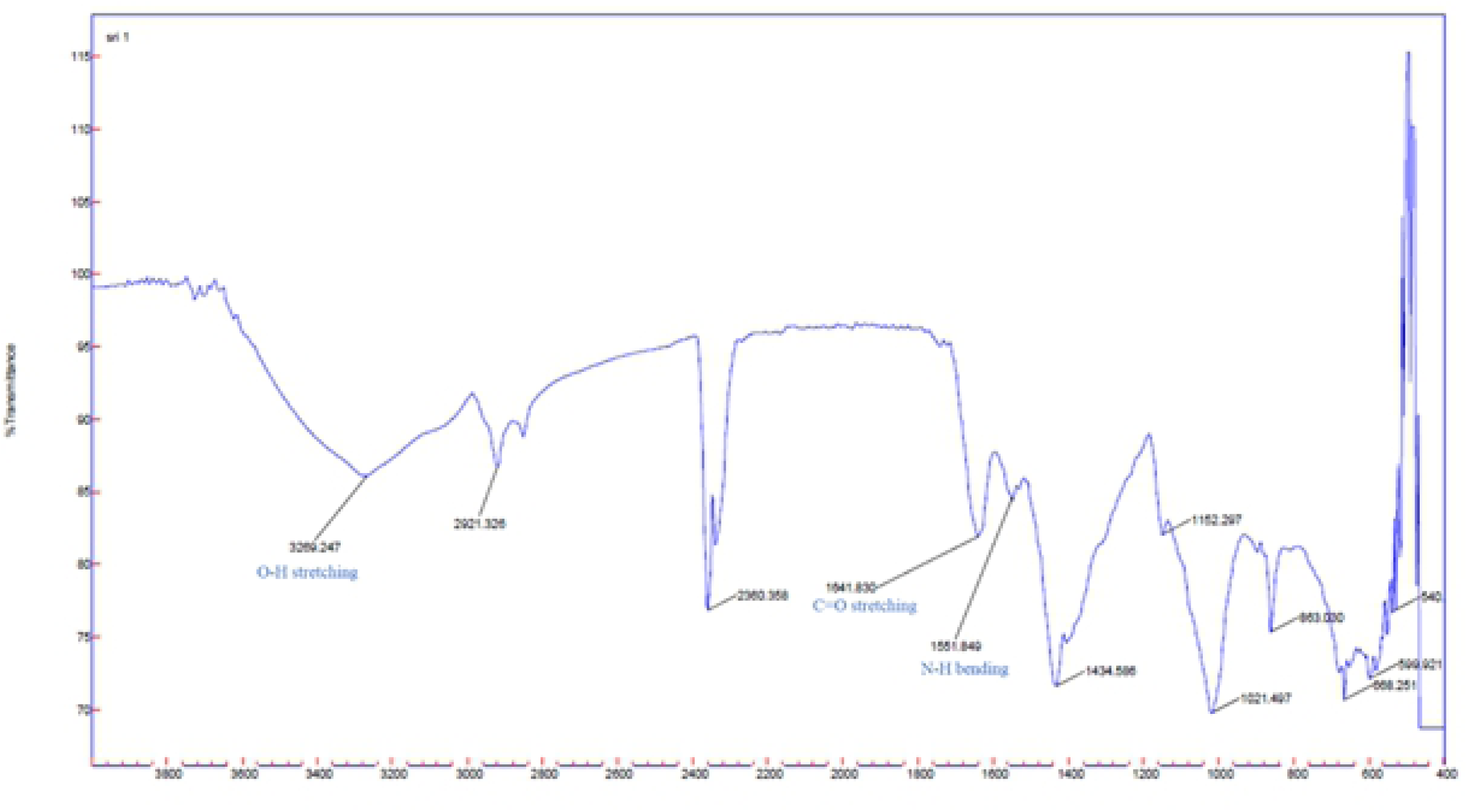
Fourier transform infrared spectra of ECM extracted *Diutina* biofilm.

### Ultrastructure of biofilm

#### Confocal scanning laser microscopy

Confocal scanning laser microscopy (CSLM) was used to visualize *D. rugosa* biofilm at 1.5, 6, 12, 24, 48, and 72 h (Fig 4). In this study, one *D. rugosa* isolate, Dr1 and one reference strain, *D. rugosa* ATCC10571 were tested. Cylindrical Intravacuolar Structures (CIVS) were produced in metabolically active yeast cells and exhibited a striking red fluorescence. FUN 1 stained the nucleic acid of *D. rugosa* cells, producing diffuse green to green-yellow cytoplasmic staining in live and membrane-compromised dead yeast cells. Dead yeast cells fluoresce with bright yellow-green with no discernible red structure.

**Fig 4.**
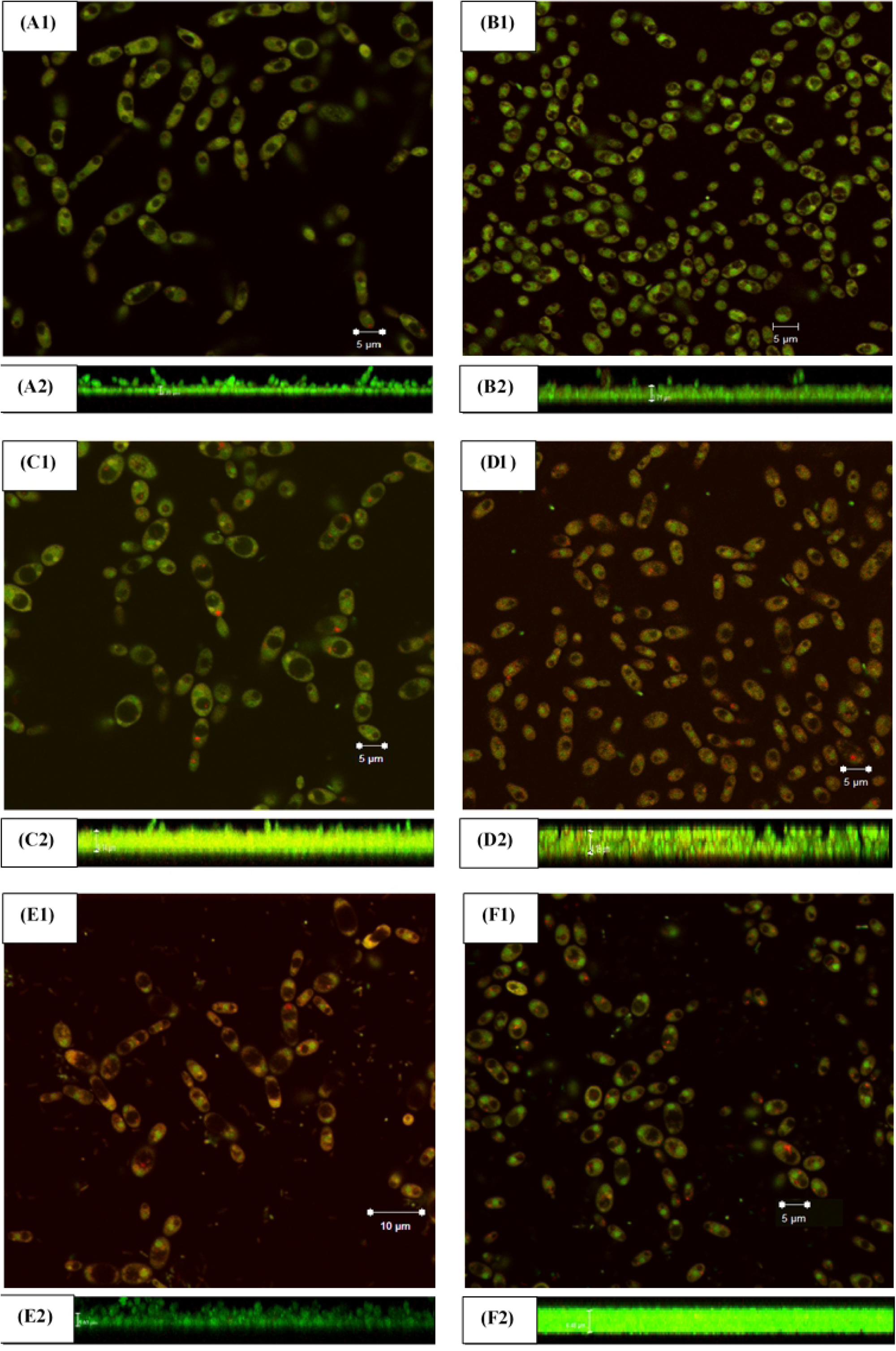
Confocal Laser Scanning micrographs of *D. rugosa* biofilm.

From the study, by 6 h, *D. rugosa* cells marked increased in density as compared to 1.5 h (adhesion phase) (Fig 4A & 4B). By 12h, microcolonies of yeast cells were visible (Fig 4C). At 24 h, the early stage of biofilm contains fungal cells are not embedded within the extracellular materials (Fig 4D). At 48h, the cell density increased with yeast cells clustered in ECM. This is observed from the regions with fluorescence (Fig 4E). The *D. rugosa* cells increased in density at 72 h (Fig 4F). Apart from the high cell density, most of the yeast cells were metabolically active yeast cells with striking red fluorescence from 12 h to 72 h.

The *D. rugosa* biofilm were grown on plastic coverslip (Thermanox, Thermo Fisher Scientific) in RPMI 1640 media at 37°C at six time points Micrographs, A1, B1, C1, D1, E1 and F1 represent the top view of *D. rugosa* biofilm and the Z-stack images on the lower side of each set are represented by A2, B2, C2, D2, E2 and F2 respectively, at various time points reveal the organization of the biofilm development. Biofilms were captured at 1.5 h (A), 6h (B), 12h (C), 24h (D), 48h (E) and 72h (F) at 100x oil immersion magnification. Arrows (white) show the formation of CIVS in metabolically active yeast cells. Arrow (dotted) show the striking red fluorescence in metabolically active cells.

Fig 5 shows the thickness and representative 3D composite images of *D. rugosa* biofilm at 1.5, 6, 12, 24, 48, and 72 h. The thickness of *D. rugosa* biofilm increases from 1.5 h to 72 h. The average thickness of the biofilm 1.5, 6, 12, 24, 48, and 72 h are 3.24 ± 0.93, 3.35 ± 1.02, 3.51 ± 1.32, 3.80 ± 1.02, 3.94 ± 0.18 and 4.50 ± 0.59 µm, respectively. The average thickness of the biofilm was measured based on three Z-stack images per sample. The Z-stack images were measured in 5 µm thick with 12 sections. During the adherence phase at 1.5 h, the yeast cells start to adhere to the plastic surface. At 6 h, the yeast cells aggregate vigorously and the cell density of the yeast cells adhered to the plastic surface increased. During the initiation phase at 12 h, the yeast cells continue to proliferate to a basal layer of biofilm on the plastic surface. The yeast cells were discernible with buddings. The early phase of maturation at 24 h, the yeast cells start to form clusters and embed in the extracellular matrix. At 48 h, the cell density of yeast cells continued to increase with yeast cells encapsulated in an extracellular matrix. During the dispersion phase at 72 h, the cell density continues to increase. However, the yeast cells appear with more yeast cells than buddings as previously observed at 12, 24, 48 h.

**Fig 5.**
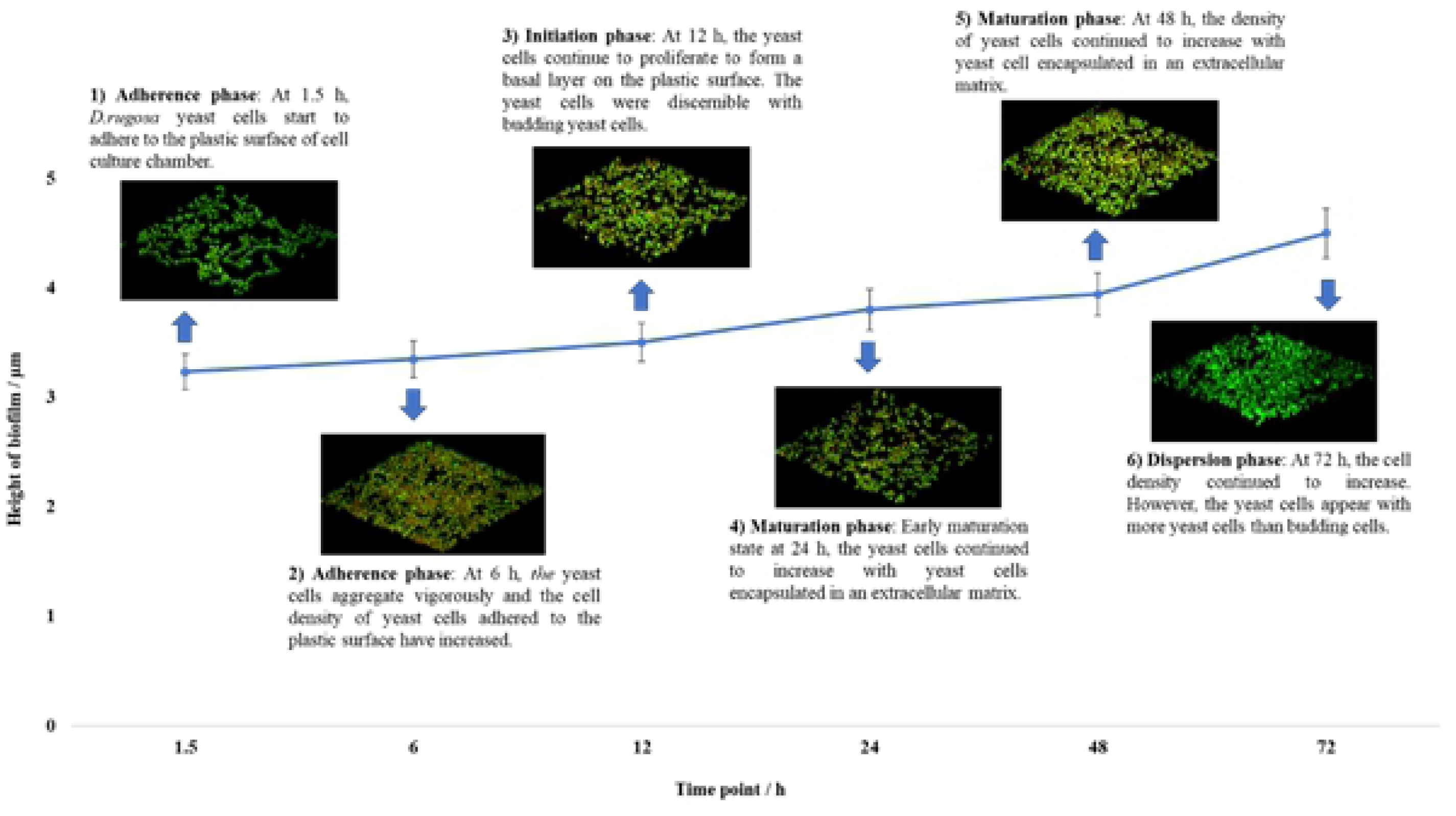
The thickness and representative 3D composite images of *D. rugosa* biofilm.

#### Scanning Electron Microscopy

SEM was used to visualize morphological changes and elucidate the biological features associated with *D. rugosa* biofilm at six time points – 1.5, 6, 12, 24, 48 and 72 h. In this study, one *D. rugosa* isolate, Dr1 and one reference strain, *D. rugosa* ATCC10571 were observed. Fig 6 shows the scanning electron micrographs of *D. rugosa* biofilm at 48 h. The *D. rugosa* biofilm was highly heterogeneous, composed of predominantly yeasts cells, blastopores and pseudohyphae. However, the yeast cells appear in a slightly elongated structure as compared to the typical round yeast morphology. The red arrows from Fig 6 shows that the *D. rugosa* cells embedded in an extracellular matrix.

**Fig 6.**
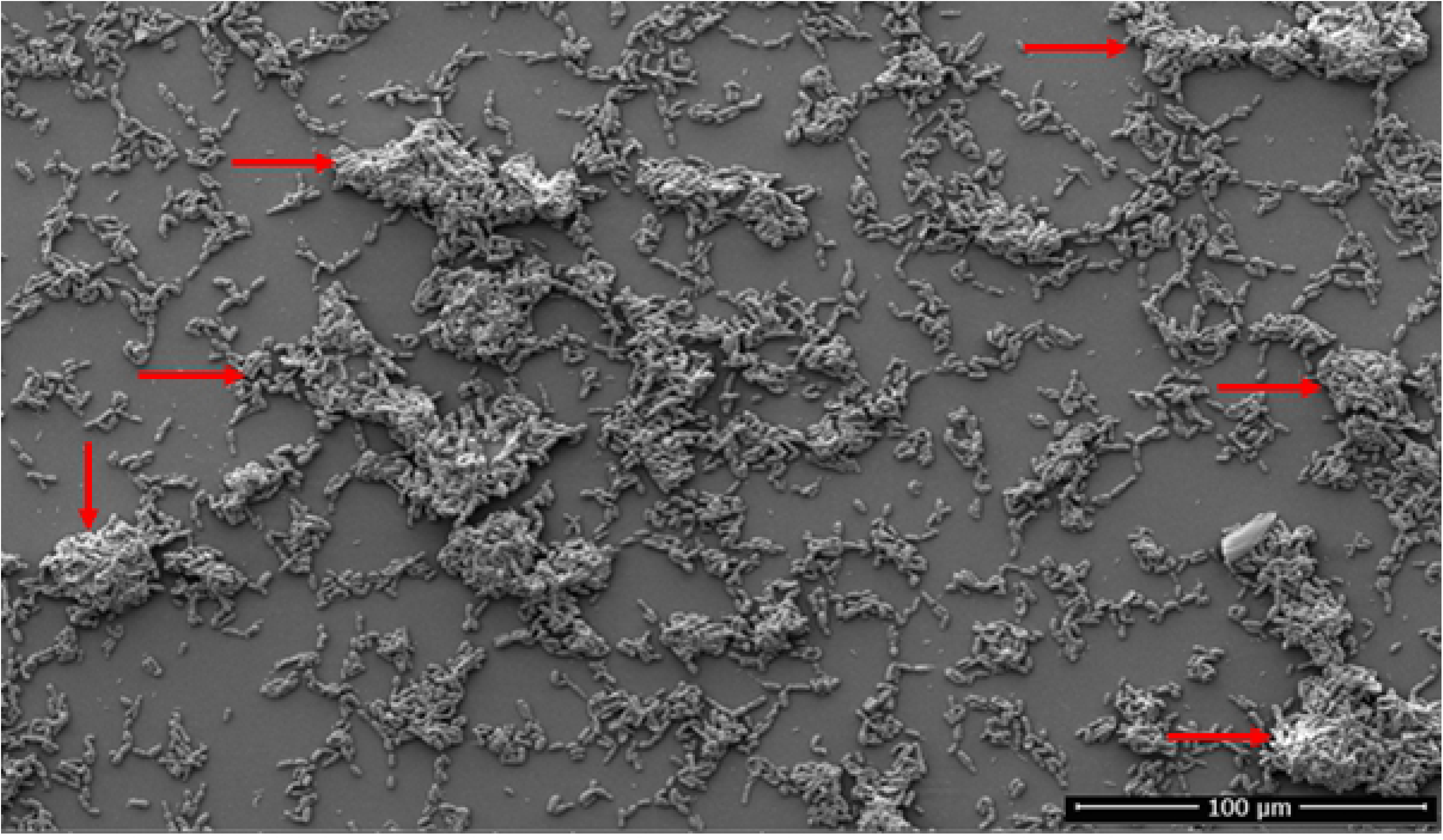
SEM micrographs of 48 h *D. rugosa* biofilm.

The arrow (red) points the *D. rugosa* cells embedded in an extracellular matrix. Magnification 1000x.

The ultrastructure of *D. rugosa* of biofilm shows that the biofilm is composed of yeast cells and pseudohyphae adhered to the plastic surface (Fig 7). In general, *D. rugosa* biofilm forms a thin biofilm that consists of blastospores embedded in an extracellular matrix and pseudohyphae. SEM analysis during the 1.5 h, adhesion phase, elongated pseudohyphae with cellular budding were observed (Fig 7A). The elongated pseudohyphae have constrictions at the position of septa. During 6 and 12 h, adhesion and initiation phase respectively, the biofilm presented with blastopores and cellular budding (Fig 7B & 7C). At 24 and 48 h, maturation phase, oval blastoconidial mother cells, pseudohyphae, sparse yeast forms and profuse budding yeast cells that forms a network were observed in *D. rugosa* biofilm (Fig 7D & & 7E). The SEM micrographs at 72 h, dispersion phase shows that the biofilm was disintegrated, with cellular budding and scar (Fig 7F). The buddings of the *D. rugosa* biofilm were mostly diploid buddings. There were no substantial differences in the ultrastructure of the biofilm between the tested isolates. Similar to CLSM findings, the cell density of *D. rugosa* biofilm increased from 1.5 h to 72 h.

**Fig 7.**
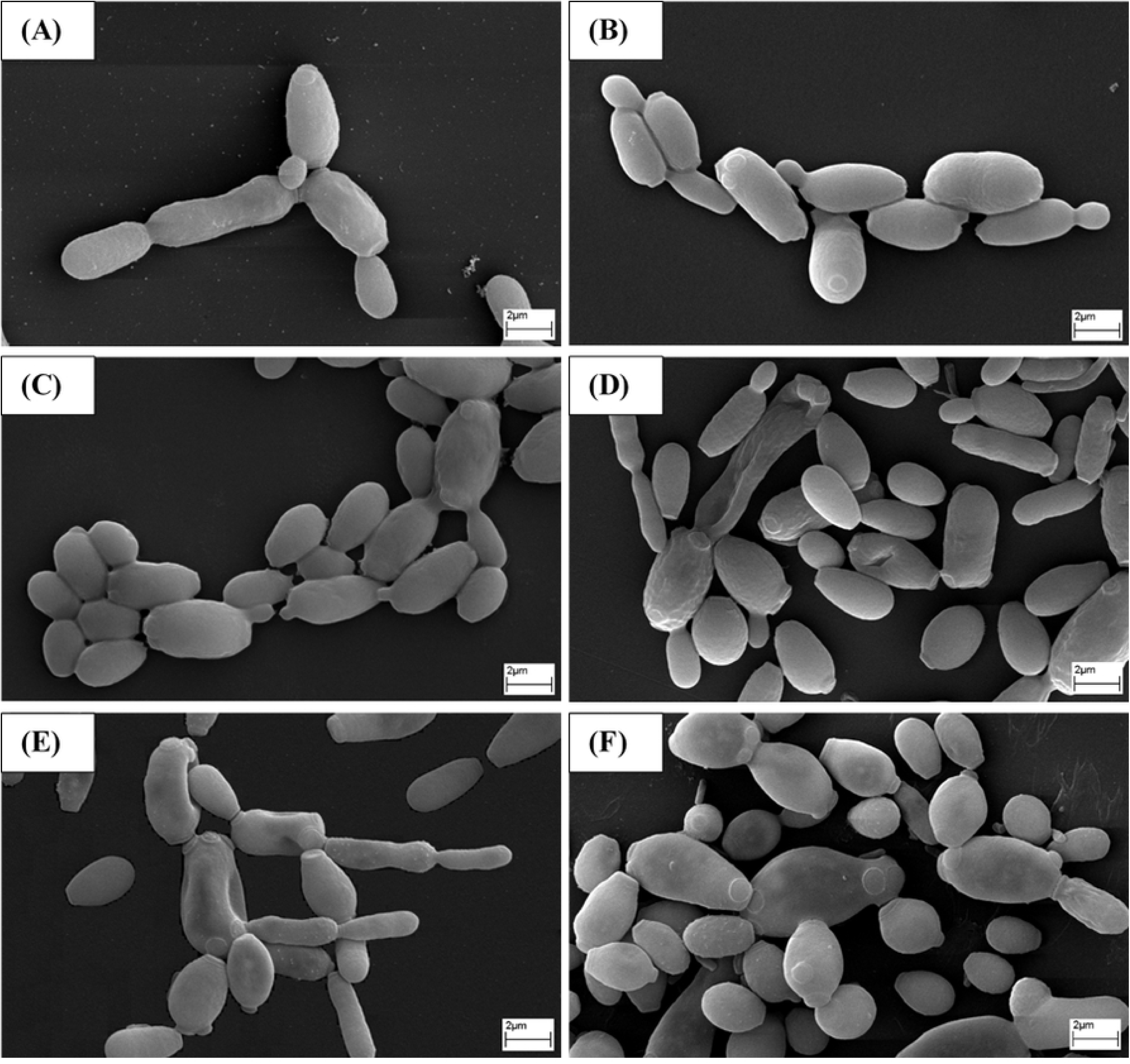
SEM micrographs of *D. rugosa* biofilm.

The *D. rugosa* biofilm were grown on plastic coverslip (Thermanox, Thermo Fisher Scientific) in RPMI 1640 media at 37°C at six time. Biofilms were captured at 1.5 h (A), 6h (B), 12h (C), 24h (D), 48h (E) and 72h (F) at 4000x magnification. Arrows (white) show the constrictions at the positions of septa.

#### Sessile Minimum Inhibitory Concentration (SMIC) of biofilm

SMICs are defined as the lowest concentration that completely inhibited the biofilm growth. The biofilm of *D. rugosa* and *D. mesorugosa* were grown on flat bottomed 96-well microtiter plates in RPMI 1640 media for 48 h prior to antifungal exposure. The antifungal susceptibility test was performed using serial doubling dilution. Fig 8 shows the graphs of antifungal susceptibility testing against *Diutina* biofilm. The values were expressed as average percentage of treated biofilm compared to untreated biofilm, calculated from calorimetric readings for XTT-reduction assays. The SMIC_50_s for the *Diutina* complex biofilm was 1024 µg/mL for amphotericin B, 512 to 2048 µg/mL for caspofungin, 256 to 2048 µg/mL for fluconazole and 256 to 2048 µg/mL for voriconazole. The SMIC 50 represents 50% reductions in metabolic activity of the biofilm. *Diutina* biofilm indicate an elevated SMICs as compared to *C. albicans* and *C. parapsilosis*. As observed in Fig 8, *Diutina* complex biofilm displayed consistent and similar values for antifungal susceptibility test, no statistically significant difference between the *Diutina* complex biofilm for amphotericin B was observed (t-test, p = 0.131). However, from antifungal susceptibility test of *Diutina* complex (*D. rugosa* and *D. mesorugosa*) on biofilms, results for caspofungin, fluconazole and voriconazole were statistically significant (t-test, p < 0.05 respectively).

**Fig 8.**
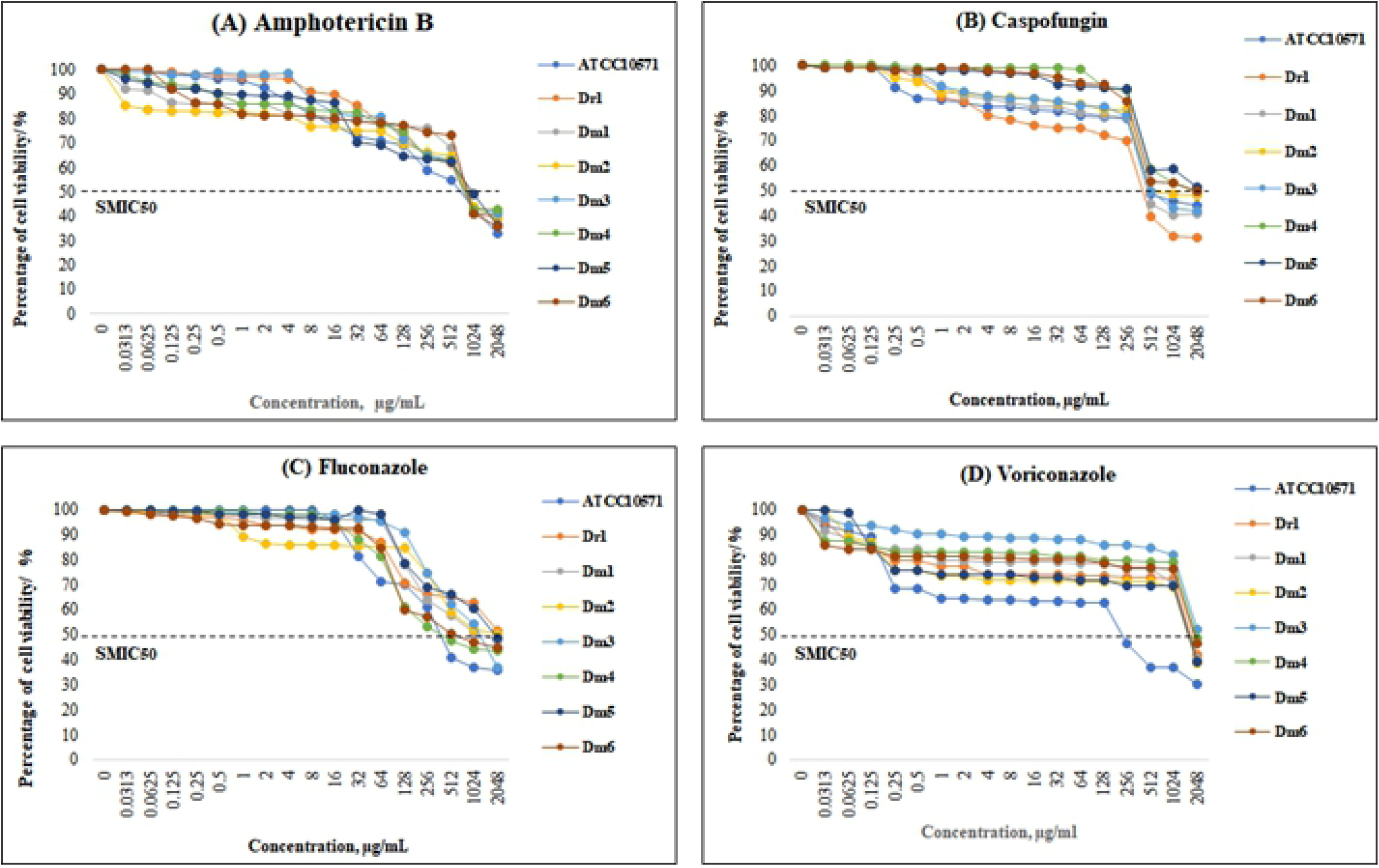
Antifungal activity of amphotericin B, caspofungin, fluconazole and voriconazole against *D. rugosa* complex biofilm.

The efficacy of different concentrations against biofilm of *D. rugosa complex*, (A) Amphotericin B, (B) Caspofungin, (C) Fluconazole and (D) Voriconazole. Values are expressed as average percentage of cell viability of treated biofilm compared to the untreated biofilm (control). The results are were expressed as mean ± standard deviations of triplicate readings and are representative of three separate experiments. SMIC_50_ is indicated by dotted lines in all graphs. Error bars represents standard deviation (n=3).

#### Minimum Inhibitory Concentration (MIC) of planktonic cells

MICs are defined as the lowest concentration that completely inhibited growth. The MIC_50_ of *Diutina* complex planktonics cells were 32 µg/mL for amphotericin B, 8 µg/mL for caspofungin, 64 to 128 µg/mL for fluconazole and 256 to 2048 µg/mL voriconazole. Fig 9 shows the graphs for the antifungal susceptibility tests against *Diutina* complex planktonic cells. The values were expressed as average percentage of treated planktonics cells compared to untreated cells. However, the MICs of *D. rugosa* complex planktonic cells for fluconazole and voriconazole were statistically significant (t-test, p < 0.05 respectively), whereas for amphotericin B and caspofungin there were no statistically significant difference (t-test, p = 0.135 and p = 0.120). The MICs for the suggested reference strains in the CLSI guideline, i.e. *C. albicans* ATCC90028 and *C. parapsilosis* ATCC 22019 were in the respective ranges in all procedures.

**Fig 9.**
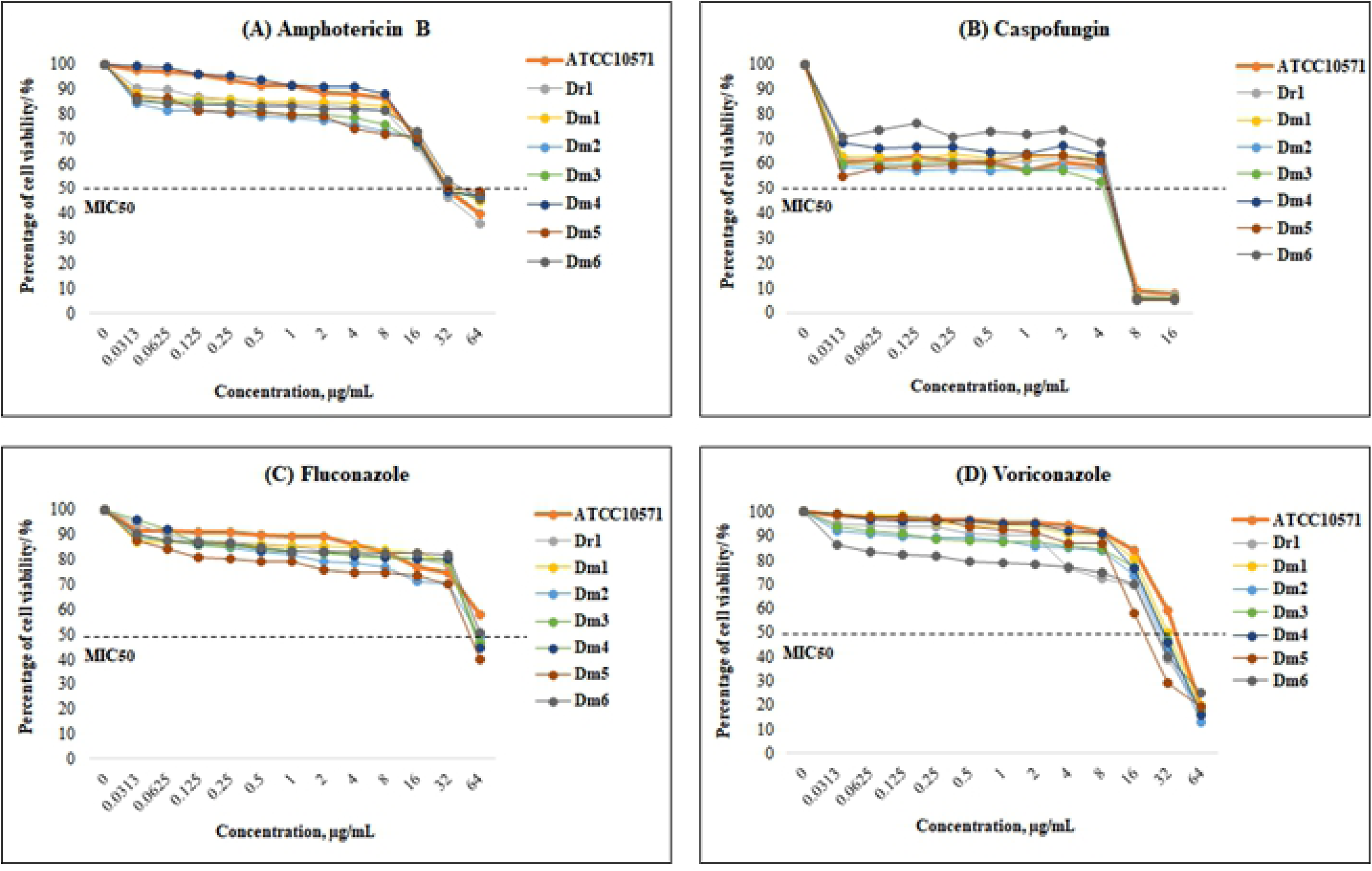
Antifungal activity of amphotericin B, caspofungin, fluconazole and voriconazole against *D. rugosa* complex planktonic cells.

The efficacy of different concentrations against *D. rugosa*, (A) Amphotericin B, (B) Caspofungin, (C) Fluconazole and (D) Voriconazole. Values are expressed as average percentage of cell viability of treated planktonic cells compared to the untreated planktonic cells (control). The results are were expressed as mean ± standard deviations of triplicate readings and are representative of three separate experiments. MIC_50_ is indicated by dotted line in all graphs. Error bars, standard deviation (n=3).

#### Penetration of antifungal agents into biofilms

In this study, CLSM is used to visualize morphological changes after exposure to antifungal agents; amphotericin B, caspofungin, fluconazole and voriconazole. SMIC_50_ from the antifungal susceptibility test using XTT assay for the antifungal agents were used to test against 48 h *D. rugosa* biofilm. Fig 10 shows CLSM micrographs of top view of *D. rugosa* biofilm and Z-stack images (lower side of each set, respectively) of preformed 48 h *D. rugosa* biofilm exposed to antifungal agents at SMIC_50_ for another 48 h and stained with Live/Dead Yeast Viability kit with FUN 1. The antifungal exposed biofilm exhibits bright, diffuse, green-yellow fluorescence that were not seen in the control. There was reduction in the thickness of the biofilm in the antifungal exposed biofilm (Fig 10).

**Fig 10.**
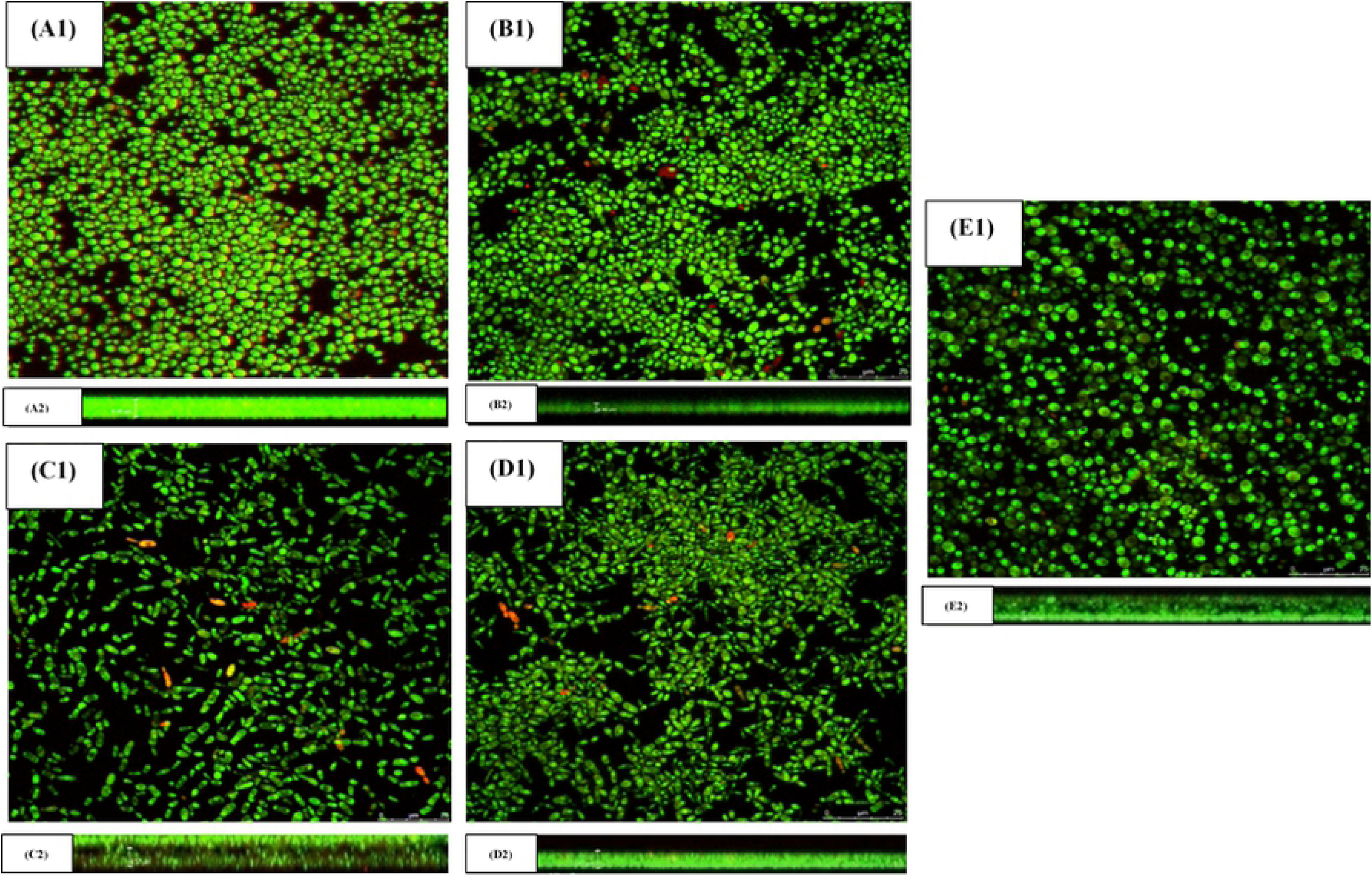
CLSM micrographs of the top view of *D. rugosa* biofilm and Z-stack images.

In concordance with the SMIC_50_ results, at 48 h, a reduced number of blastospores and shrunken vacuoles were observed in biofilm exposed to 1024 µg/ml of amphotericin B, 512 µg/ml of caspofungin, 2048 µg/mL of fluconazole and 2048 of µg/ml voriconazole as shown in Fig 10. The red fluorescence CIVS and large vacuole observed in the control was not seen in treated biofilm.

CLSM micrographs of the top view of *D. rugosa* biofilm in control (without any antifungal treatment) (A1) and cells treated with 1024 µg/mL amphotericin B (B1), 512 µg/mL caspofungin (C1), 2048 µg/mL fluconazole (D1), 2048 µg/mL voriconazole (E1) at 100x oil immersion magnification. The Z-stack images of the are represented by A2, B2, C2 and D2 at the lower side of each set. Biofilm were grown on polystyrene surface in a 2-well chamber slides in RPMI 1640 media for 48 h at 37°C and then exposed antifungal agents for another 48 h. *D. rugosa* biofilm were stained with Live/Dead Yeast Viability kit with FUN 1.

Data from the Z-stack images used to measure the height of the biofilm with and without (growth control) exposure to antifungal agents. Interestingly, the Z-stack images that show the height of *D. rugosa* biofilm exposed to all four antifungal agents decreased as compared to the control *D. rugosa* biofilm, as shown in Fig 11. The Z-stack images were measured in 5 µm thick with 12 sections.

**Fig 11.**
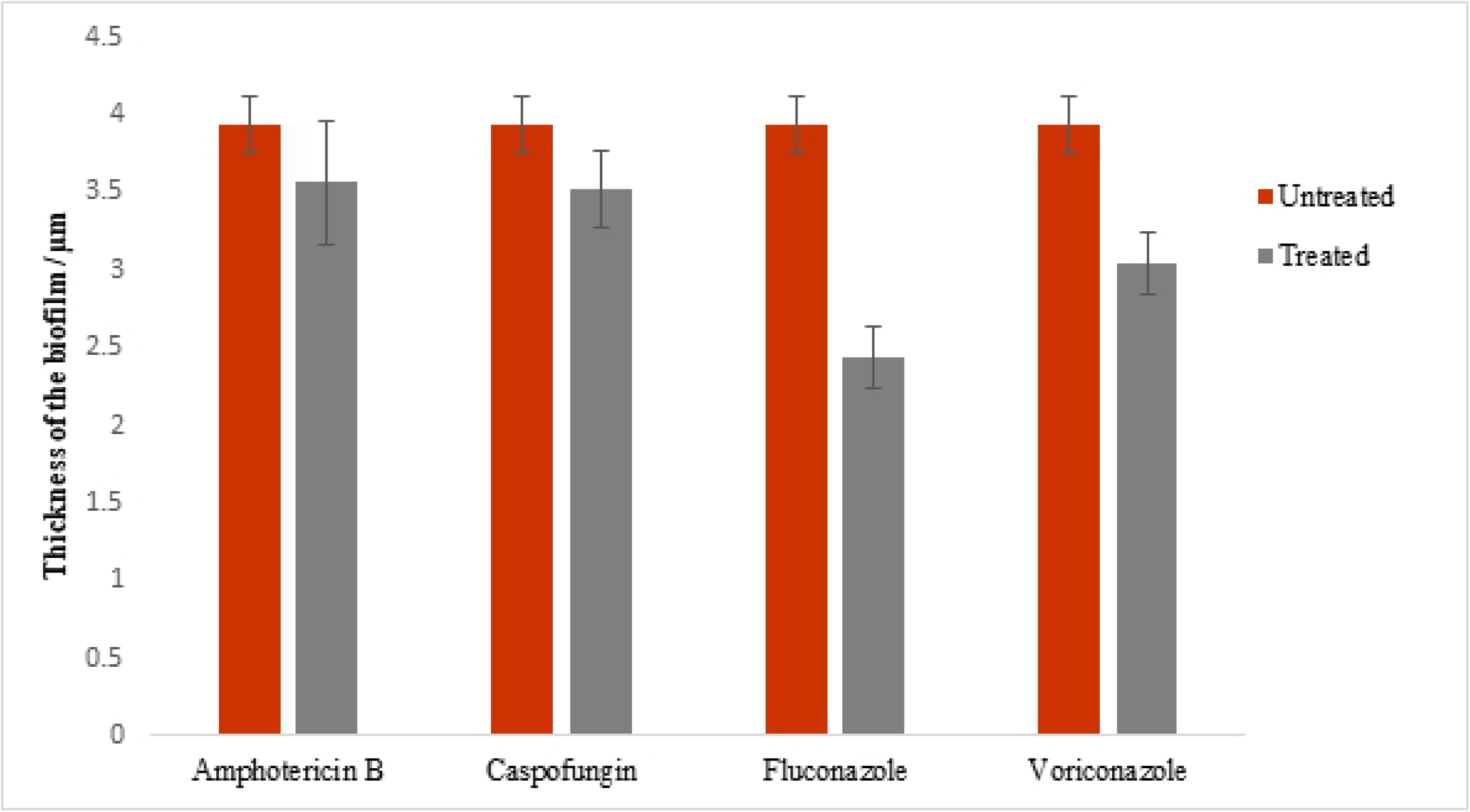
The thickness of *D. rugosa* biofilm pre and post exposure to antifungal agents.

The thickness of 48 h D. rugosa biofilm after exposure to antifungal agents at SMIC_50_ (grey bars) and *D. rugosa* biofilm without exposure to antifungal agents (control) (red bars). The results are were expressed as mean ± standard deviations of triplicate readings and are representative of three separate experiments. Error bars, standard deviation (n=3).

The SEM micrographs were used to observe the microstructure of *D. rugosa* post-exposure to antifungal agents, i.e. amphotericin B, caspofungin, fluconazole and voriconazole (Fig 12). In this study, SMIC_50_ from the antifungal susceptibility test using XTT assay for the antifungal agents were tested on 48 h *D. rugosa* biofilm.

**Fig 12.**
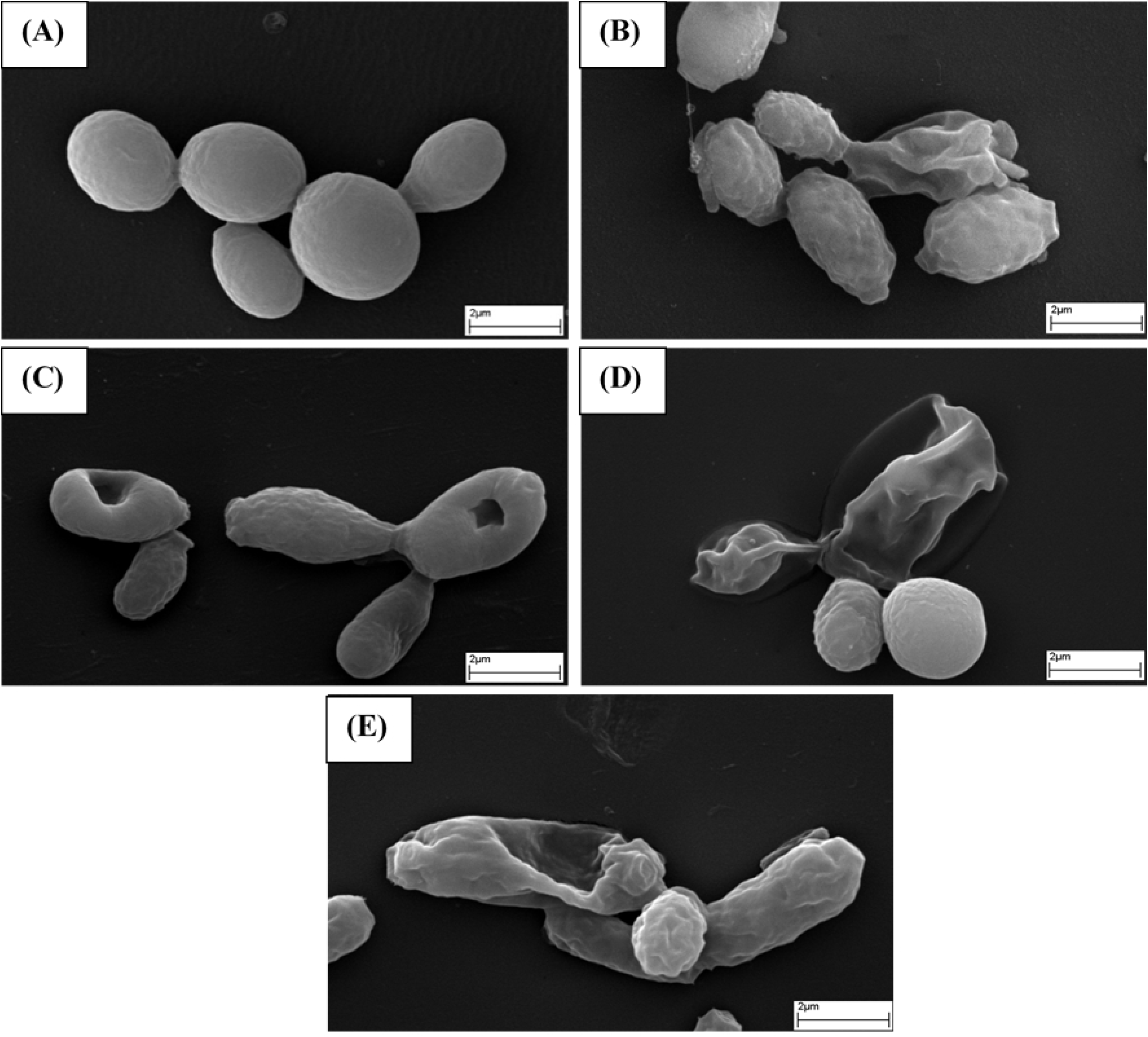
SEM micrographs of D. *rugosa* biofilm pre and post exposure to antifungal agents.

SEM micrographs of preformed *D. rugosa* (A) without antifungal drugs and in the presence of antifungal agents; (B) Amphotericin B, 1024 µg/mL (C) Caspofungin, 512 µg/mL (D) Fluconazole, 2048 µg/mL and (E) Voriconazole, 2048 µg/mL. All micrographs were captured at 8000x magnification.

Exposure to antifungal agents shows a reduction in the fungal biomass and altered yeast morphology. The structure of the yeast cells collapsed and was disfigured when exposed to amphotericin B (Figure 12B), fluconazole (Figure 12D) and voriconazole (Figure 12E). On the other hand, exposure to caspofungin (Figure 12C) shows the presence of punctured yeast morphology. Figure 12A shows the control, *D. rugosa* biofilm without antifungal drugs treatment.

## Discussion

Little is known about the biofilm-forming ability of the newly reclassified genus *D. rugosa* complex and to our knowledge, this is the first study on biofilm ultrastructure of *D. rugosa* complex. In a previous study, Seneviratne and co-workers have explained that the virulence of *Candida* species correlates with biofilm formation and its architecture rather than biofilm biomass production [31, 32]. The variation in the biofilm extracellular matrix may explain the differences in metabolic activities between species [33]. Biofilm forming ability is highly depending on the type of species. The z-stack images from CLSM micrographs revealed that the thickness of *D. rugosa* biofilm increased from a period between 1.5 h to 72 h. The drawback of the thickness measurement using z-stack images is the inability to distinguish on live/dead cells. As such, the thickness measurement is inclusive of live and dead cells adhered to the abiotic surface. Further visualization of biofilm ultrastructure by SEM revealed that *D. rugosa* is a poor biofilm producer with less dense yeast network and pseudohypha elements. Mature *D. rugosa* biofilm (24 h - 48 h) is composed of a basal layer of blastospores covered by sparse yeast cells with pseudohypha embedded in extracellular matrix. Here, branched pseudohyphal filament (Fig 13A) and elongated tube-shaped pseudohyphal filament (Fig 13B) were observed predominantly during the maturation stage. The pseudohyphae formed branched and elongated blastoconidial cells with constrictions at the septal junction where septal junctions are classic representation of pseudohyphal forms [34]. Sónia Silva and colleagues have reported that *C. albicans, C. parapsilosis* and *C. tropicalis* are true polymorphic yeasts, whereas *C. albicans* and *C. tropicalis* are capable to grow as blastospores, hyphae and/or pseudohyphae. However, *C. parapsilosis* commonly grows as blastospores, and, it can sometimes produce pseudohyphae [35]. In contrary, *C. glabrata* is not a polymorphic organism, growing as blastospores only [25, 35]. The ability of *Candida* species to switch from yeast to hyphal and pseudohyphal is strongly associated with its virulence, both the former and latter structures assist in tissue penetration during *Candida* infections and plays a vital role in organ metastasizing [36]. From this study, according to the SEM micrographs, *D. rugosa* is a polymorphic yeast, although, blastospore growth is prevalent at all time points, pseudohyphae are generated in the early stages of biofilm formation (Fig 13).

**Fig 13.**
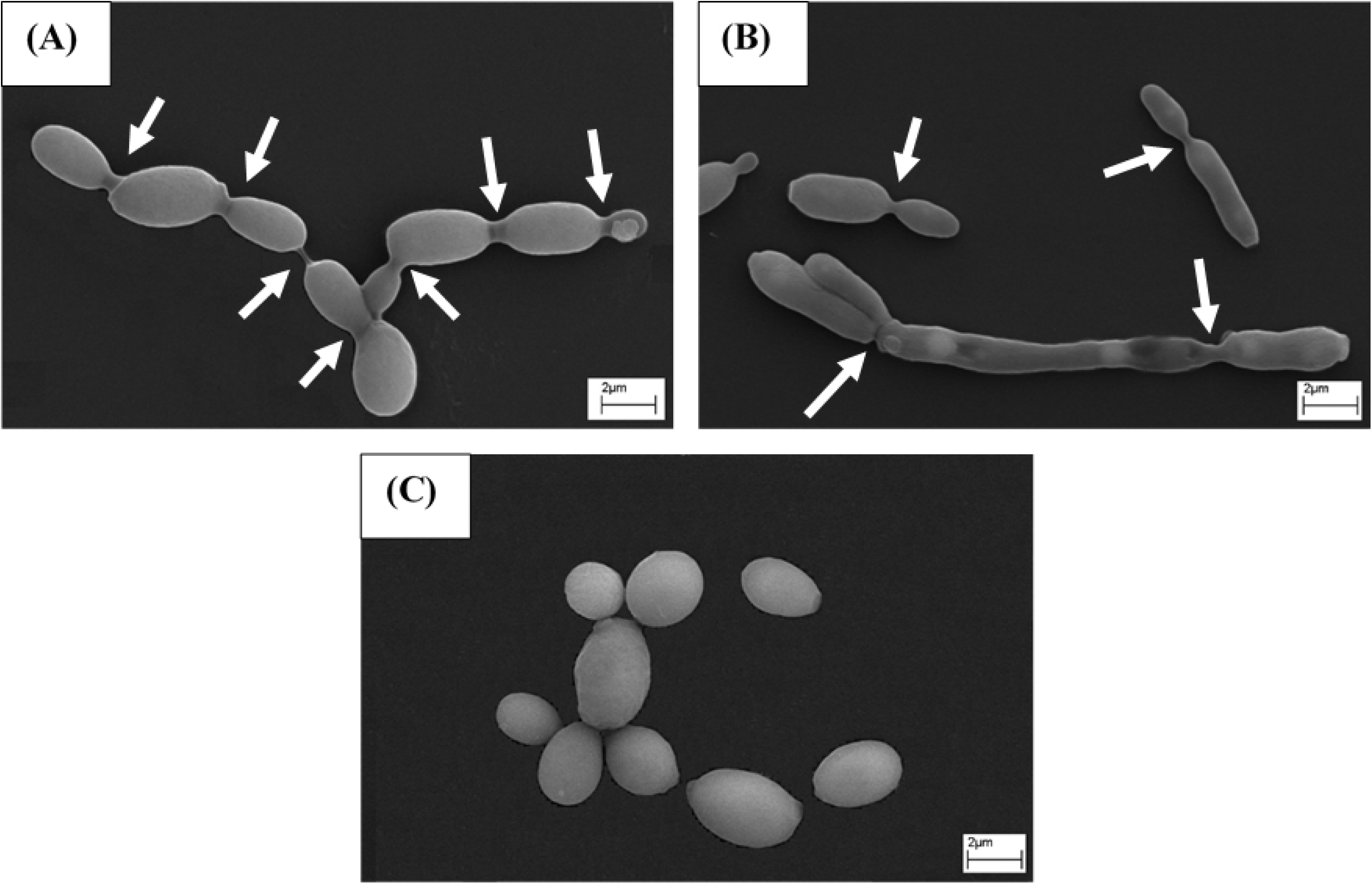
SEM micrographs of *D. rugosa* biofilm morphologies.

*D. rugosa* biofilm grown on plastic coverslip – yeast and pseudohyphae. Yeast cells showing branched pseudohyphal filament with bipolar buddings in which daughter cells are formed at both poles of a pseudohypha (A), elongated pseudohyphal filament with distal budding (B) and non-dividing yeast cells (C). All micrographs were captured at 4000x magnification. Arrows point to constrictions at septal sites.

Biofilm formed on abiotic surfaces have substantial similarities to biofilm formed on living tissue (biotic) such as mucosal layers [37]. Shantanu Ganguly and colleague have reported that some of the major genetic determinants are similar in biofilm found in vaginal candidiasis and denture biofilm model [37]. Nevertheless, *in vitro* and *in vivo C. albicans* biofilm models have presented with distinct morphology and characteristics [37-41]. In *in vitro* model, the basal layer of *C. albicans* biofilm are primarily composed of yeast cells with filamentous elements and the ECM is rich in beta-glucan [37-39, 42]. In contrary, in an *in vivo* model, *C. albicans* biofilm presented in more disorderly manner with interspersed yeast cells with filamentous element and ECM that contains host immune cells such as neutrophils [37, 40, 41]]. The present study focuses on *in vitro* biofilm formation on abiotic surface that will provide a foundation to embark on *in vivo* studies.

*D. rugosa* outbreak have been reported globally. Data on its epidemiology suggests that *D. rugosa* was most commonly isolated from bloodstream and urinary tract infections [4, 5, 7]. Despite, Kocyigit I et al. have reported first continuous ambulatory peritoneal dialysis (CAPD) related peritonitis caused by *D. rugosa* [43]. Furthermore, the high prevalent of *D. rugosa* infections in intravenous catheter could be associated to biofilm formation of *Candida* species. Biofilm are notable to be resistant to antifungal agents up to 1000-fold more than their planktonic counter parts [15, 16]. The particular interest of this study was to assess the resistance/susceptibility of *D. rugosa* biofilm towards antifungal agents specifically to amphotericin B, caspofungin, fluconazole and voriconazole. Resistance to antifungal agents has heighten in *Candida* species, particularly in non-albicans species. Till date, there are four drug classes available for systemic *Candida* infection treatment including the azoles (fluconazole, itraconazole, isavuconazole, posaconazole and voriconazole), polyenes (amphotericin B), echinocandins (anidulafungin, caspofungin, and micafungin) and lastly, the pyrimidine (flycytosine) [44]. Even so, only azoles, polyenes and echinocandins are licensed for monotherapy for *Candida* infections and only fluconazole and echinocandins are proposed as first-line antifungal agents for invasive candidiasis [44].

Many studies have reported that *D. rugosa* isolates have decreased susceptibility to fluconazole [45-47]. Similarly, *D. rugosa* are notable for decreased susceptibility to amphotericin B [48]. Amphotericin B belongs to polyene group antifungal agent and is the antifungal agent of choice for the treatment of systemic fungal infections. Amphotericin B disrupts the fungal cell wall synthesis by binding to ergosterol which lead to the formation of pores that allow leakage of cellular components. Amphotericin B is well known for its toxicity. On the contrary, the new class of antifungal agents such as echinocandin works by inhibiting the enzyme, 1,3-β-D-glucan synthase hence disrupting the integrity of fungal cell wall. The echinocandins are being widely used as therapy for invasive candidiasis particularly for *Candida* species that are resistant to polyenes and azoles groups of drugs. There are clinical guidelines that recommend echinocandins as first-line therapy for candidemia and infection caused by *C. parapsilosis* [49-51]. Nonetheless, there have been evidence of echinocandin resistance in patients with *Candida* infection including *C. albicans, C. glabrata, C. krusei and C. parapsilosis* [52-56].

The findings of this study showed that biofilm of *D. rugosa* complex isolates have elevated SMICs for all the four antifungal agents tested (Table 1). Similarly, *D. rugosa* planktonic cells have elevated MICs for amphotericin B, fluconazole and voriconazole (Table 1). However, Chunyan Ming and co-workers have reported that *D. rugosa* complex (*D. rugosa, D. mesorugosa, D. pseudorugosa* and *D. neorugosa*) are sensitive to amphotericin B and fluconazole (MIC 0.5 – 1 µg/mL) [57]. Interestingly, *D. rugosa* complex planktonic cells are more susceptible to caspofungin as compared to amphotericin B, fluconazole and voriconazole. The abundance of 1,3-β-D-glucan in *D. rugosa* cell wall may attribute to its susceptibility to caspofungin. The interpretive breakpoint of antifungal agents for *D. rugosa* planktonic cells have not been established in CLSI or EUCAST guideline. Although, the results from this study suggest that *D. rugosa* complex may be multi drug resistant species, the low number of isolates tested in this study is insufficient to support the suggestion.

**Table 1.** Antifungal susceptibility data for *D. rugosa* biofilm and planktonic cells.

| Strains | Amphotericin B |  | Caspofungin |  | Fluconazole |  | Voriconazole |  |
| --- | --- | --- | --- | --- | --- | --- | --- | --- |
|  | SMIC 50 ± SD | MIC 50 ± SD | SMIC 50 ± SD | MIC 50 ± SD | SMIC 50 ± SD | MIC 50 ± SD | SMIC 50 ± SD | MIC 50 ± SD |
| *D. rugosa*<br>ATCC 10571 | 1024 ± 0.001 | 32 ± 0.021 | 512 ± 0.001 | 8 ± 0.004 | 512 ± 0.006 | 128 ± 0.085 | 256 ± 0.001 | 64 ± 0.047 |
| Dr1 | 1024 ± 0.002 | 32 ± 0.039 | 512 ± 0.001 | 8 ± 0.006 | 2048 ± 0.001 | 64 ± 0.08 | 2048 ± 0.001 | 32 ± 0.086 |
| Dm1 | 1024 ± 0.001 | 32 ± 0.034 | 1024 ± 0.001 | 8 ± 0.009 | 1024 ± 0.004 | 64 ± 0.148 | 2048 ± 0.001 | 32 ± 0.017 |
| Dm2 | 1024 ± 0.001 | 32 ± 0.015 | 2048 ± 0.001 | 8 ± 0.008 | 1024 ± 0.004 | 64 ± 0.061 | 2048 ± 0.001 | 32 ± 0.062 |
| Dm3 | 1024 ± 0.004 | 32 ± 0.023 | 2048 ± 0.001 | 8 ± 0.009 | 1024 ± 0.004 | 64 ± 0.148 | 2048 ± 0.001 | 32 ± 0.024 |
| Dm4 | 1024 ± 0.001 | 32 ± 0.018 | 512 ± 0.003 | 8 ± 0.003 | 256 ± 0.001 | 64 ± 0.147 | 2048 ± 0.001 | 32 ± 0.033 |
| Dm5 | 1024 ± 0.004 | 32 ± 0.039 | 2048 ± 0.002 | 8 ± 0.006 | 2048 ± 0.001 | 64 ± 0.052 | 2048 ± 0.001 | 32 ± 0.027 |
| Dm6 | 1024 ± 0.007 | 32 ± 0.052 | 512 ± 0.001 | 8 ± 0.005 | 512 ± 0.002 | 64 ± 0.117 | 2048 ± 0.001 | 32 ± 0.010 |

SMIC, sessile minimum inhibitory concentration; MIC, minimum inhibitory concentration, SD, standard deviation.

The lifestyle of *Candida* species biofilm is defined by the constituents of the ECM, a unique acquisition acquired by *Candida* species biofilm that provides structural integrity, mediates adhesive and cohesive interactions, contributes mechanical stability and supply nutrients for the consortium cells [58-59]. Pioneer researchers have mentioned that the ECM of *C. albicans* are composed of four major macromolecular classes, including polysaccharides, proteins, lipids and nucleic acids [59, 60]]. Previous studies have reported that ECM from *C. albicans* biofilm consist of carbohydrate, protein, phosphorus and hexosamine [59]. Every *Candida* species biofilm have distinctive amount of protein and carbohydrate in their ECM; however, the carbohydrate fraction exhibits highest degree of heterogenicity within *Candida* species biofilm [59]. *C. glabrata* and *C. tropicalis* have high protein content with low carbohydrate content, in contrary, *C. parapsilosis* have high carbohydrate content with low protein content [61] Similarly, in this study, *D. rugosa* biofilm exhibited with high carbohydrate content with low protein content. Nevertheless, the ECM may be influenced by environmental factors including temperature, pH, oxygen, growth medium, etc [59]. Previous studies have reported that increased matrix production was observed in *C. albicans* biofilm grown under flow condition compared to static condition [62, 63]. Extraction of ECM of *Candida* species can be challenging, and it is important to ensure that the extraction procedure specifically removes matrix and does not extract cell wall component. In this study, extraction procedure was carried out using sonication and purification according to the pioneering studies with slight modifications [64, 65].

Perhaps the most primordial role of ECM is its role as physical barrier for antifungal agent penetration, although, antifungal resistance is multifactorial [59]. Several studies have now implicated that β-1,3-glucans is a putative component of ECM in *Candida* species that play a part in antifungal sequestration particularly to azole and polyene group of drugs where binding to β-1,3-glucans prevents the antifungal agents reaching their cellular target [40]. In addition, it is notable that echinocandins such as caspofungin binds to β-1,3-glucan synthase to block the synthesis of β-1,3-glucans and causing disruption of the fungal cell wall integrity and eventually leads to cell death. β-1,3-glucans are essential components of *Candida* cell wall and elevated β-1,3-glucans were observed in the ECM of biofilm both *in vitro* and *in vivo* [62]. A study by Nett J et al. revealed that biofilm treated with β-1,3-glucanase are more susceptible to fluconazole [40]. In this study, β-glucan were detected in the cell wall of *D. rugosa* complex (2.5-4.2%), however, β-glucan were not detected in the ECM of *D. rugosa* complex biofilm. The ECM extraction method could be one of the reasons for the undetected β-glucan. This study uses sonication method for ECM extraction, alternative approach such as mechanical (glass beads) extraction method can be used to solve the problem. The yield of β-glucan was higher in the *D. rugosa* complex isolates than the *D. rugosa* ATCC 10571. This result corroborates with previous finding of β-glucan yield in *D. rugosa* [66]. More in depth analyses using a combination of biochemical approaches and analytical chemistry leading to identification of the components of ECM in *D. rugosa* biofilm is required, particularly the constituents of carbohydrate that facilitate the understanding of its antifungal resistance traits.

## Conclusion

The findings of the study provide information on the ultrastructure, antifungal susceptibility and biochemical analysis of the biofilm matrix of *Diutina* complex, known to be a rare pathogenic yeast. In assessing the sensitivity of the biofilm to antifungal agents, all isolates studied were susceptible to high concentration of amphotericin B, caspofungin, fluconazole and voriconazole. However, caspofungin displayed a potent *in vitro* activity against the planktonic cells. Findings from CLSM and SEM confirmed changes in the biofilm morphologies upon exposure to antifungal agents at the their respective SMICs. The study further demonstrates that the biofilm matrix was mainly composed of carbohydrates with small amount of proteins, without any β-glucan content. However, β-glucan were only detected in the cell wall of planktonic cells of *D. rugosa* complex.

## Acknowledgements

This study was supported by Ministry of Education Malaysia (FRGS/2/2014/SKK01/TAYLOR/02/1) and Taylor’s University (TU) for funding and technical support.

